# Standing covariation between genomic and epigenomic patterns as source for natural selection in wild strawberry plants

**DOI:** 10.1101/2021.03.31.437859

**Authors:** Hanne De Kort, Tuomas Toivainen, Filip Van Nieuwerburgh, Bart Panis, Timo P. Hytönen, Olivier Honnay

## Abstract

Adaptive evolution is generally thought to be the result of natural selection predominantly acting upon pre-existing DNA sequence polymorphisms through gene-environment interactions. Epigenetic inheritance is, however, recently considered an additional molecular force driving adaptive evolution independent of DNA sequence variation. Through comparative analyses of genome-wide genetic (SNPs) and epigenetic (DMCs) variation of wild strawberry plants raised under distinct drought settings, we demonstrate intermediate levels of genome-wide covariation between SNPs and DMCs. Cases of high SNP-DMC covariation were significantly associated with (i) applied stress, (ii) non-adaptive SNPs, and (iii) solitary DMCs (as opposed to DMC islands). We also found that DMCs, drought-responsive DMCs in particular, typically co-vary with hundreds of SNPs, indicating high genomic redundancy as a basis for polygenic adaptation. Our findings suggest that stress-responsive DMCs initially co-vary with many associated SNPs under increased environmental stress (cfr. co-gradient plasticity), and that natural selection acting upon these SNPs subsequently reduce standing covariation with stress-responsive DMCs. In addition, the degree of covariation between SNPs and DMCs appears independent of their respective genomic distance, indicating that trans-acting associations between SNPs and DMCs are as likely as cis-acting associations. Our study is in favor of DNA methylation profiles representing complex quantitative traits rather than independent evolutionary forces, but further research is required to fully rule out SNP-independence of genome-wide DMCs. We provide a conceptual framework for polygenic regulation and adaptation shaping genome-wide methylation patterns.

## Introduction

The ability of natural populations to adapt to novel environmental stressors is vital to biodiversity, particularly in an era of accelerated environmental change. Understanding the molecular mechanisms that allow populations to withstand environmental perturbations therefore is fundamental to biodiversity conservation. Quantitative genetics theory predicts that the probability of populations to keep pace with environmental change increases with molecular variation underlying phenotypic traits affecting fitness^1–3^. While heritable genetic variation is thought to be the dominant driver of trait evolution, a rising number of studies suggests an important role for epigenetic variation and non-genetic inheritance in governing adaptive trait-environment covariation^4–6^. It remains elusive, however, to what extent genetic variation dictates epigenetic responses to environmental changes^7^. If genetic variation drives all genome-wide epigenetic and regulatory processes in response to environmental stress, then epigenetic variation should be considered a consequential quantitative trait that is under genetic control, rather than an independent evolutionary force.

Most adaptive epigenetic variation may thus, directly or moderated by environmental influences, result from genetic variation, questioning the importance of epigenetic adaptation as an ubiquitous independent evolutionary mechanism. Uncoupling epigenetic from genetic evolution is, however, challenging, requiring experimental work on isogenic or inbred lines each raised under various environmental settings and preferably over multiple generations, or comparative studies integrating genomic and epigenomic analysis of genotypes that underwent distinct evolutionary trajectories. Experimental inbred studies show that environmental stress generates ample epigenetic variation^8–10^, but that epigenetic responses to treatments are genotype-specific^9,10^ (but see^11^). The observation that each inbred line generates a specific epigenetic profile may suggest non-independence between DNA sequence polymorphisms and differential methylation^9,10^. Comparative genome-epigenome studies, on the other hand, find many more differentially methylated genomic regions than DNA sequence polymorphisms, and little evidence for causative mutations underlying epigenetic variation (but see^12,13^, suggesting that (i) part of the epigenetic variation can arise independent from genetic evolution^14,15^, and/or (ii) single DNA sequence polymorphisms (e.g. in methyltransferase enzymes catalysing genome-wide DNA methylation) pleiotropically affect epigenetic variation across large genomic regions (e.g.^16^K). Molecular mechanisms such as polygenic and pleiotropic regulation of epigenetic variation likely are omnipresent throughout the genome of most species^17–20^, posing an extreme challenge to disentangling epigenetic from genetic adaptive evolution.

Multivariate tools capturing the various aspects of gene-epigene co-variation may help to assess the overall tendency of epigenetic variation to coincide with genetic variation, and to identify epigenetic signatures that cannot be explained by genetic variation. Studies comparing genetic and epigenetic structure in field-collected individuals frequently find indications of cryptic, environment-driven epigenetic population structure not coinciding with genetic variation^4,5,21,22^. This is, however, not surprising considering that gene expression and associated epigenetic variation are the result of gene×environment interactions allowing the same genetic background to render distinct phenotypes. Controlling for environmental effects is thus crucial for identifying epigenetic signatures that can be transmitted to the next generation independent of genetic inheritance (see e.g.^23^).

DNA methylation islands, i.e. genomic regions featured by high-density methylation, can have large epigenetic effects on gene expression and phenotypic fitness^24–26^. For example, up to 90% of heritability for flowering time could be explained by a small number of islands of differential methylation in isogenic *Arabidopsis* lines^24^. Many DNA methylation islands are, however, linked to DNA sequence variation in transposable elements, gene bodies and intergenic regulatory elements^27–29^. Genomic islands of differential methylation, i.e. DNA methylation islands with significantly different methylation levels between populations, may therefore particularly arise near genetic polymorphisms that are under the influence of divergent natural selection (see also^30^). Identifying to what extent islands of differential methylation can arise independent of genetic variation may help assessing the role of genome-wide methylation landscapes in adaptive evolution.

Epigenetic variation may contribute disproportionally to adaptive evolution where the amount of standing genetic diversity is low, for example in clonal species, following demographic bottlenecks, and/or where selection pressures are variable or extremely high^31–37^. A recent study comparing population epigenomic signatures between a steep altitudinal gradient and a much wider continental gradient in the clonal species *Fragaria vesca* (woodland strawberry), demonstrated that genomic islands of epigenetic divergence readily arise even at fine spatial scale (<2km), and persist through the next generation^6^. How these epigenetic patterns behave relative to genetic variation nevertheless remains an open question that is key to understanding the molecular basis of adaptive evolution.

Aiming to shed new light on the interplay between epigenetic and genetic adaptive evolution, we compared genome-wide methylation profiles with genome-wide SNP (single nucleotide polymorphism) data from a leaf of 20 *F. vesca* individuals originating from three nearby populations and raised in a common garden to control for unstable environment-driven epigenetic signatures. We used a hierarchical sampling design involving a steep altitudinal population gradient, which has been demonstrated to generate fine-scale adaptive trait divergence in *F. vesca*^38^, and subjected half of the individuals of one population (N=10) to acute drought stress. This treatment helped disentangling DMC patterns that are expected to be gene-independent (stress-induced) from SNP-associated DMC patterns. Through distinguishing between neutral and adaptive SNPs (genetic outliers) we also tested whether differential methylation between natural populations is associated to adaptive evolution acting upon nearby SNPs. We then explored to what extent acute stress, adaptation evolution, genomic proximity, and the tendency of DMCs to co-occur in genomic islands of differential methylation together explain genome-wide epigenetic and genetic co-variation. Gene ontology enrichment tests served to assess whether SNP-DMC covariation is reflected by specific biological functions. We finally tested to what extent genes associated to flowering time, an important fitness trait with high heritability and adaptive divergence in *F. vesca*^38^, are under genetic and/or epigenetic control. Flowering genes, which regulate vernalisation and timing of flowering in response to various environmental stimuli including light conditions and temperature, are prime examples of adaptive DNA sequences that can be subject to evolutionary divergence (e.g.^39^) yet of which transcription is finetuned by epigenetic modifications (e.g.^24,40^).

## Methods

### Study system

*Fragaria vesca* is a self-compatible clonal species of deciduous forest edges and gaps, with a relatively small diploid genome (2n = 14; 240 Mbp^41^). Seeds were collected from *F. vesca* plants at three locations along a fine-scale altitudinal gradient in the French Pyrenees (<2kms between populations, coordinates), including five plants from a low and high altitudinal population (400 and 900 masl, resp.), and ten plants from a mid-altitudinal population (600 masl). Clear adaptive trait divergence has previously been observed along the fine-scale gradient (covering 9 populations, with ca. 13 plants per population), coinciding with various topographical variables including elevation, aspect and slope (De Kort et al. 2020). Among the most heritable and adaptive traits was flowering vigour per unit biomass, demonstrating that this fitness trait is under strong selection even at a very fine spatial scale.

After germination in petri-dishes, one random seedling per plant (N=20) was grown in a growth chamber with standardized soil moisture and light conditions. To help disentangling genetic from epigenetic variation, five samples from the mid Pyrenees were raised under drought stress through applying three consecutive drought treatments starting two months after germination. At the start of each treatment, watering stopped until leaves went limp (6-10 days), after which plants were watered for three days to allow partial recovery of the soil. The plants were allowed to rehydrate for one week after the third drought treatment to remove most instable drought-induced epigenetic effects. The plants were kept at room temperature in a growth chamber with regular growth lamps. Ca. three months after germination, one leaf per plant was collected in liquid nitrogen prior to DNA extraction.

### Whole-genome bisulfite sequencing and methylation profiling

Library preparation, bisulfite conversion and whole-genome sequencing occurred following De Kort et al. 2020^6^. Briefly, paired-end sequencing was performed on 30 whole genome DNA fragment libraries obtained after DNA extraction and sonication, rendering 76 417 704 ±13 837 490 high quality reads (mean ±standard deviation) per sample of 75 bp. One sample from Poland (“East”) showed an increased level of duplicate reads and was excluded from further analysis. Average sequencing depth after mapping and deduplication was 30x.

Methylkit v1.10.0^42^ was used to call significant differentially methylated cytosines (DMCs), which were deemed significant where (i) at least 3 samples per population had a minimum cytosine coverage of 5x, (ii) at least a 25% difference in methylation between populations was observed, and (iii) q-values <0.01. DMCs were called (see De Kort et al. 2020^6^ to demonstrate that alternative DMC calling parameters do not considerably affect methylation patterns). We adopted a missing data threshold of 6.9% (max. 2 out of 29 data points missing). These missing data were imputed with the population-specific average methylation level (Supporting table 1). Because DMCs often tend to cluster together, we assessed the tendency of DMCs to co-occur (< 100 bp between DMCs) along genomic blocks of 1 kb using the genomic R package “bumphunter v1.12”. We further defined DMC islands where at least five DMCs were found within these 1 kb blocks. Cytosines with significantly differentiated methylation levels between the three populations and between the drought stress treatment are referred to as inherited and drought-responsive DMCs, respectively.

### Outlier analysis

We filtered our SNP dataset to only keep polymorphic SNPs with at least two individuals harbouring the minor allele (corresponding to an average MAF of 0.04). The remaining SNPs that were physically linked (<100 bp along linkage blocks of 1 kb) were thinned using the R package “bumphunter v1.12”. From the linked SNPs, we selected the SNP with least missing data. We further allowed one missing data point per population (average missing data rate of 13.7%), and imputed these missing data with the population-specific average genotype. This rendered a total of 17,755 SNPs (Supporting table 2).

PCadapt was used to identify SNPs with exceptional genetic patterns that reflect adaptive genetic differentiation^43^. While F_ST_ outlier methods are prone to false results (i.e. neutral loci falsely identified as adaptive loci), PCadapt has been demonstrated to outperform alternative outlier detection methods through accounting for background genetic structure, particularly in the presence of hierarchical population structure^43^. In addition, our prime interest does not lie in identifying candidate SNPs of selection, but to assess whether, on overall, presumably adaptive SNPs are more important than neutral SNPs in explaining epigenetic variation. In PCadapt, we accounted for linkage disequilibrium through removing correlated SNPs that are highly correlated (r^2^>0.25) within windows of 200 base pairs while computing principal components representing background genetic structure. A total of five principal components captured most background genetic variation (K=5). The SNPs that deviated significantly from neutral background structure along the principal components (expressed as the Mahalanobis distance) were identified as putatively adaptive loci. We applied a false discovery rate cut-off of 1% to correct for multiple testing using the R package “qvalue”.

### DMC-SNP covariation

To assess covariation between epigenetic and genetic population structure, we first performed a Principal Component Analysis (PCA) on the methylation and allele frequency matrix, respectively. Individual genotypes were coded as 0, 1 or 2 representing allele counts (homozygous for reference allele, heterozygous, and homozygous for alternative allele). Then, to identify the general degree of covariation between genetic and epigenetic population structure, we performed a coinertia analysis between the principal components of the SNP matrix and the principal components of the DMC methylation percentages (R package “ade4”)^44^. This rendered 28 co-inertia axes delineating the multidimensional molecular space, of which the first two axes covered most covariation between allele and methylation frequencies (75.7%). The overall strength of the covariation between SNPs and DMCs was measured through the RV-coefficient, and its significance was determined using 1000 Monte Carlo permutations. Mahalanobis distances, representing the dissimilarity between SNPs and DMCs in the two-dimensional coinertia space, were calculated for each SNP-DMC pair, using the R package “*StatMatch*”. The degree of covariation between each SNP-DMC pair was calculated as [maximum Mahalanobis distance between SNP and DMC scores - pairwise SNP-DMC Mahalanobis distance], divided by the maximum Mahalanobis distance to obtain a covariation index between 0 (corresponding to the maximum Mahalanobis distance) and 1 (corresponding to maximum covariation). The SNP-DMC pairs with a covariation index > 0.9 were considered as co-varying SNP-DMCs (n=5,687 ~ 0.33% of all SNP-DMC pairs).

A mixed model was generated to test whether the degree of covariation between genetic and epigenetic structure depended upon (i) the logarithmic distance between each SNP and all DMC islands, (ii) natural selection (outliers vs. background SNPs), (iii) DMC clustering (number of linked DMCs in the DMC island), and (iv) acute drought stress (drought-responsive DMCs vs. inherited DMCs), while controlling for SNP and DMC cluster ID (two random factors). This model served to address the following hypotheses in respective order: (i) SNP-associated DMCs are predominantly cis-acting (i.e. close to covarying SNP) rather than trans-acting, (ii) SNP-associated methylation differentiation is linked to natural selection acting upon SNPs rather than to neutral processes, (iii) SNP-DMC covariation is more likely where more dense DMC islands are involved, and (iv) SNP-DMC covariation is more pronounced for DMCs inherited from parental environmental conditions than for drought stress-induced DMCs.

The covariation between SNPs and DMCs may not be reflected by specific alleles directly resulting into higher or lower methylation frequencies, but may rather reflect alleles altering the methylation sensitivity or *variability* of co-varying DMCs. To test how the allelic composition of SNPs alters methylation profiles of co-varying DMCs, we first calculated methylation variability as the coefficient of variation (SD/mean) in methylation frequencies across the samples. A higher coefficient of variation thus corresponds to higher methylation variability. We then tested the amount of SNP-DMC covariation explained by (i) the absolute difference in average methylation *frequencies* between SNPs that were homozygous vs. heterozygous for the reference allele (Δmethylation frequency, normalized to obtain values between 0 and 100), and (ii) the absolute difference in methylation *variability*between SNPs that were homozygous vs. heterozygous for the reference allele (Δmethylation variability). In addition, we hypothesize that the relationship between SNP-DMC covariation and methylation patterns differed between DMCs arising from drought stress than for inherited DMCs that were not affected by drought stress (fixed effect “DMC stress”), because acute environmental stress may reduce variation in methylation frequencies. Finally, because only a tiny fraction of the SNPs was homozygous for the alternative alleles (0.25%), these were excluded from the mixed model:

Covariation ~ (Δmethylation frequency ×DMC stress) + (Δmethylation variability ×DMC stress) + SNP (random) + DMC cluster (random)

## Results

A total of 1,611 SNPs (9.02%) significantly deviated from background genetic structure (Fig. 1A). Many SNPs (n=93) were within or near a total of 66 flowering genes, eight of them (10.81% of flowering genes) showing signatures of fine-scale adaptation (Fig. 1A).

**Fig. 1.**
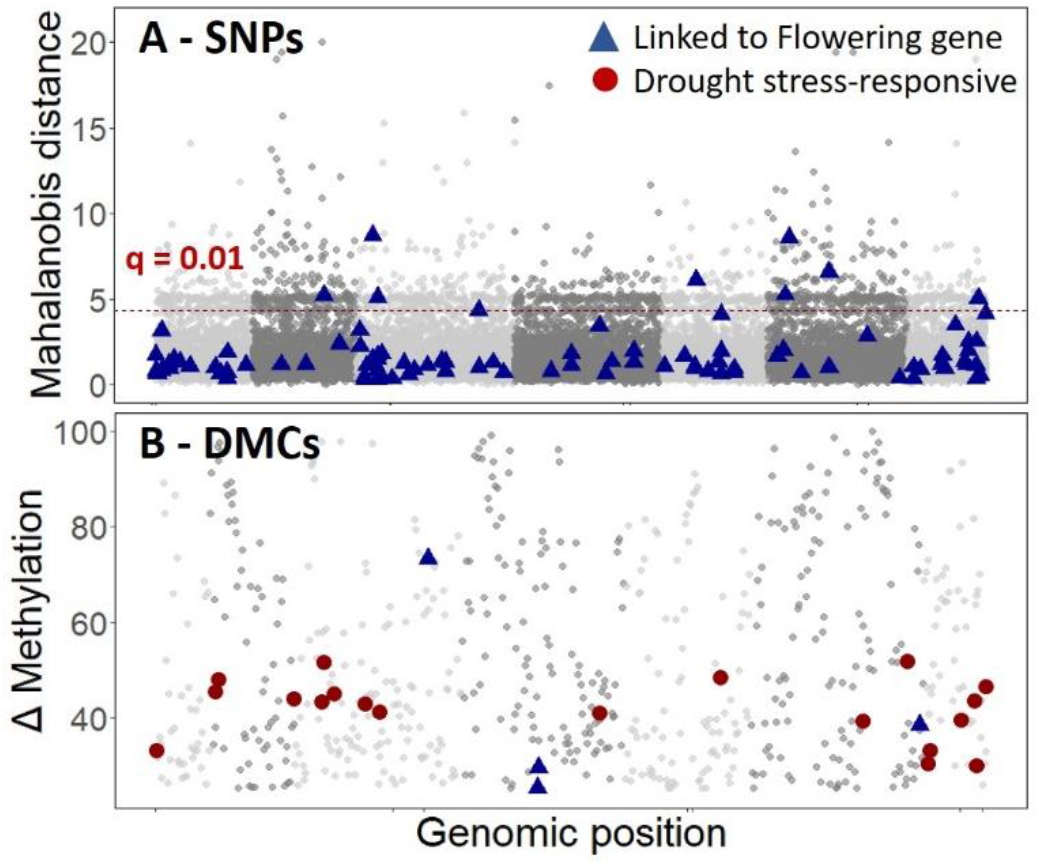
Genome-wide distribution of genetic (A) and methylation (B) polymorphisms along a fine spatial gradient. SNPs above the red dashed line are significant outliers. Alternate grey-colors represent the seven chromosomes.

A total of 619 DMCs were found along this gradient, most of which were found in solitude or loosely linked (79.16%) while 20.84% clustered together in genomic islands of differential methylation (Fig. 1B). These DMC islands represent genomically clustered DNA methylation signatures that significantly differentiated along the altitudinal gradient, and contained on average 10 DMCs (up to 29 DMCs) within 1kb linkage blocks. Drought stress gave rise to 19 solitary DMCs, and no DMC islands (Fig. 1B). Although we did not obtain methylation profiles from flowers, we found significant DMCs within or near flowering genes (Fig. 1B, Fig. 2A), suggesting their involvement in consecutive, altitude-dependent expression of flowering genes.

**Fig. 2.**
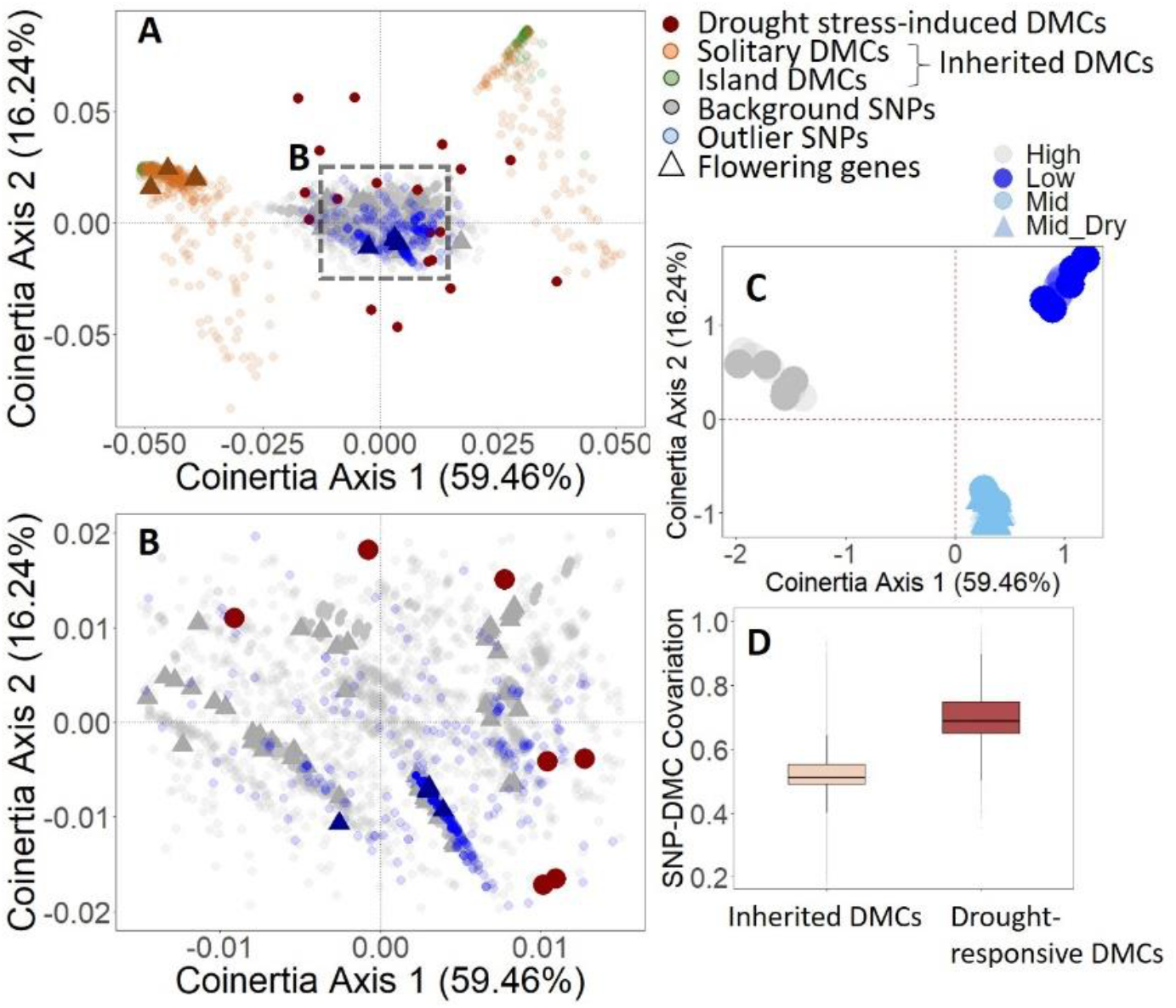
SNP-DMC covariation patterns along two coinertia axes (A-C) and between drought-responsive vs. droughtinsensitive DMCs (D).

The multivariate co-inertia analysis revealed moderate levels of covariation between genome-wide SNP and DMC signatures (coefficient of covariation = 41.97%, p<0.001, Fig. 2), and most covariation aligned with the altitudinal gradient (first axis in Fig. 2C). Although genotypes raised under drought stress clustered together with genotypes that did not receive a drought treatment (light blue genotypes in Fig. 2C), drought-responsive DMCs showed markedly high covariation with SNPs (i.e. they clustered together with the SNPs, Fig. 2A and B). A total of 5,681 SNPs (32.00%) co-varied with a total of 18 drought-responsive DMCs (2.91%), and these SNPs were significantly more likely to co-vary with drought-responsive DMCs than with inherited DMCs (Fig. 2D, Fig. 3A, Table 1).

**Fig. 3.**
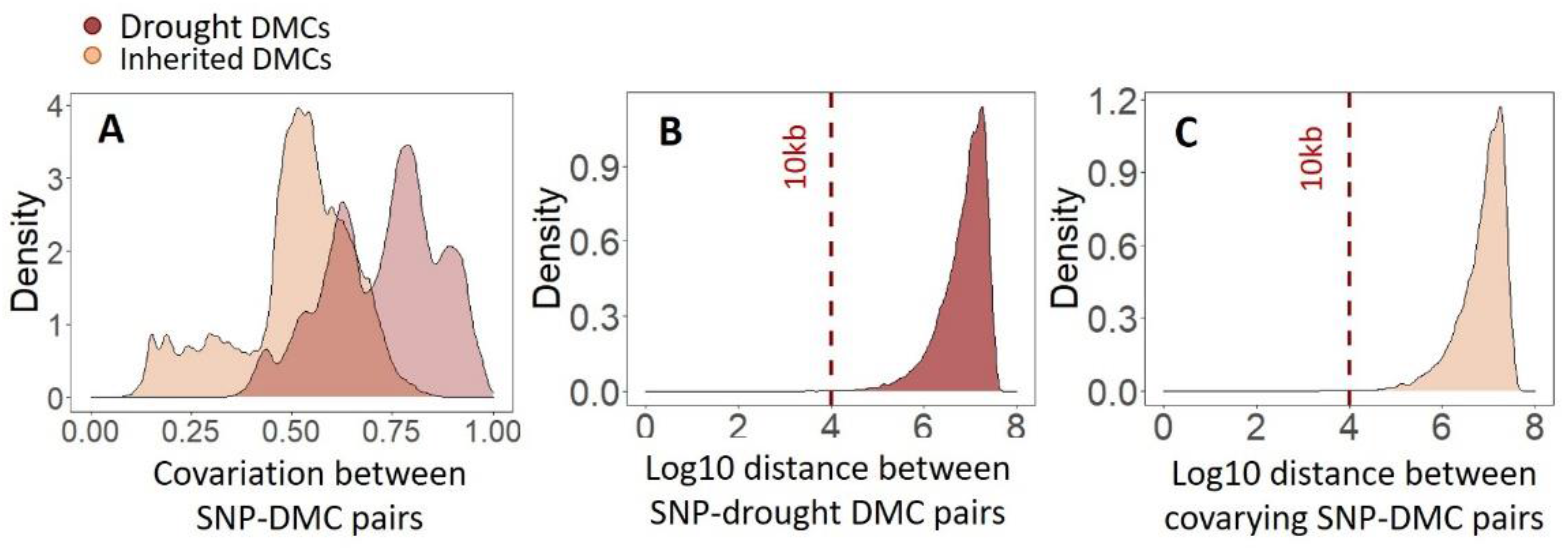
Density plots showing distribution of SNP-DMC covariation (A) and genomic distance (B,C).

**Table 1.** Mixed model results. The left model tested which SNP and DMC features explain SNP-DMC covariation. The right model tested whether high SNP-DMC covariation results in differential methylation frequency rather than differential methylation variability. The table provides ANOVA F-values with asterisks representing significance (** for p<0.01 and *** for p<0.001), and R^2^ showing marginal (fixed effects only) and conditional (full model) explained variance (%), respectively. The strongest effects are in bold.

| SNP-DMC covariation variables |  |  | DMC signature associated with SNP-DMC covariation |  |  |
| --- | --- | --- | --- | --- | --- |
| | F | $R^2$ | | F | $R^2$ |
| Genomic distance (Log10) | 7.21** | 24.31/<br>84.37 | $\Delta$ methylation frequency [1] | 0.58 | 10.07/<br>82.76 |
| **Selection** (outlier vs. backgr. SNP) | 1356*** |  | **$\Delta$ methylation variability [2]** | 851*** |  |
| DMC (drought-induc. vs inherited) | 52.31*** |  | DMC (drought vs inherited) | 60.53*** |  |
| DMC clustering (sqrt) | 17.59*** | | [1] × DMC | 41.80*** | |
| **Selection × DMC clustering** | 2312*** |  | **[2] × DMC** | 523*** |  |

Covariation between SNPs and DMCs not only was relatively high for drought stress-induced DMCs, but also depended on SNP and DMC features. Specifically, and contrary to expectations, significantly higher covariation with DMCs was found for background SNPs as opposed to outlier SNPs (Fig. 4), and covariation of DMCs with SNPs was higher for solitary DMCs than for DMC islands (Fig. 4B, Table 1).

**Fig. 4.**
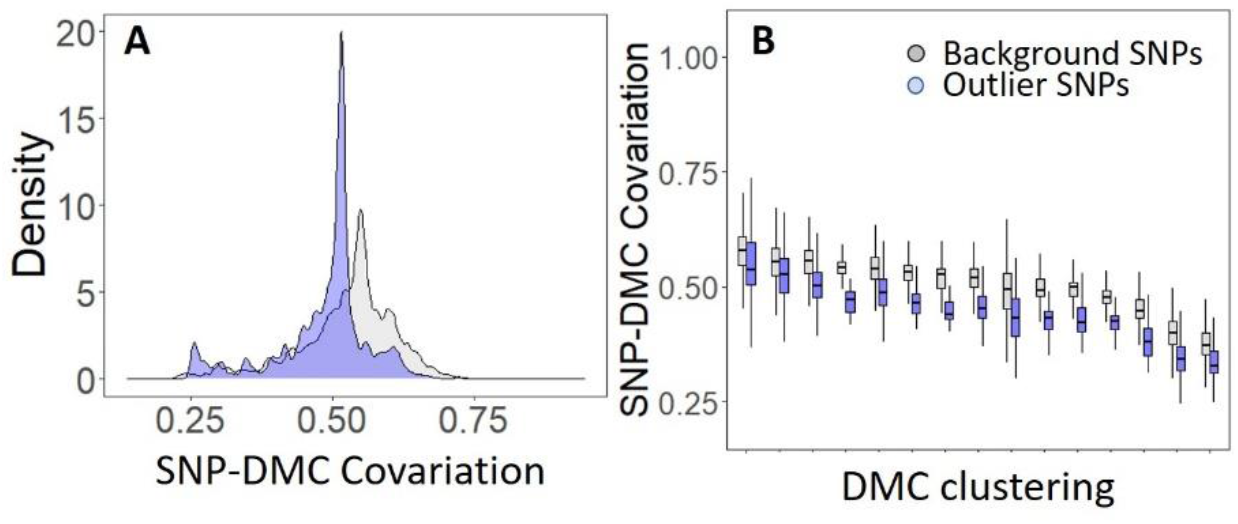
The amount of covariation between pairs of SNPs and DMCs across the genome (A) and depending on genomic DMC clustering (from solitary DMCs up to dense DMC islands) (B). Marginal covariation was used to exclude pseudoreplication of shared SNPs and DMC islands across SNP-DMC pairs.

Covariation between a DMC and a SNPs did not translate into obvious associations between allele frequencies and methylation frequencies. Instead, co-varying SNPs were associated with differential methylation variability, but only for inherited DMCs (Fig. 5, Table 1). Drought-induced DMCs, which had highest covariation indices, did not display altered methylation variability (Fig. 5), likely because drought stress canalized methylation levels. In all co-varying SNP-DMC pairs involving inherited DMCs (n=35 with a covariation index > 0.9), SNPs resulted in higher vs. lower methylation variability when they were homozygous vs. heterozygous for the reference allele.

**Fig. 5.**
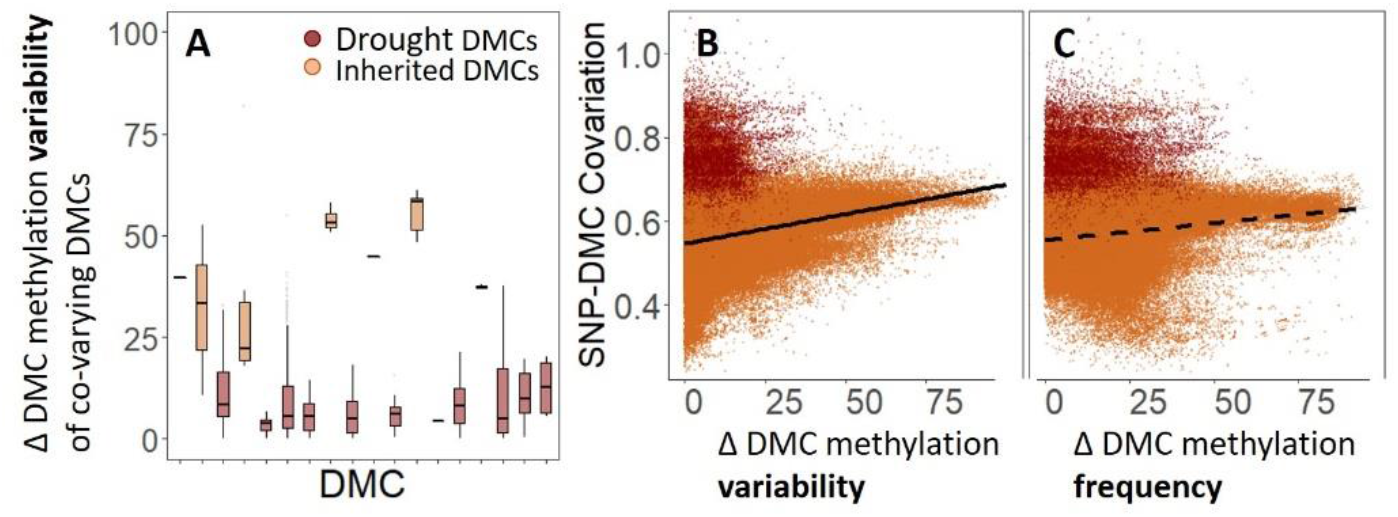
Relationships between pairwise SNP-DMC covariation and differential DMC methylation associated with the allelic composition of the respective SNP.

## Discussion

Epigenetic variation is frequently linked to environmental and fitness differences, implying its involvement in adaptive evolution. DNA sequence-independent inheritance of DNA methylation signatures is, however, much more challenging to demonstrate, particularly when DNA methylation marks can be triggered by multiple genetic polymorphisms. Such polygenic control at single base resolution dramatically complicates the identification of causal SNP-DMC associations. Here, we use multivariate techniques to demonstrate that direct SNP-DMC associations are inexistent or too hard to detect, and that SNP-DMC covariation most likely is linked to genomic redundancy. Natural selection appears to reduce SNP-DMC covariation through reducing the proportion of alleles affecting methylation at DMCs involved in fitness. All results point to differential DNA methylation marks representing quantitative traits that can be the target of selection rather than sources for geneindependent evolution. Based on our findings, we propose a framework of polygenic adaptation shaping genome-wide methylation levels.

Theoretical and empirical data have shown that adaptive evolution of quantitative traits is a highly dynamic and polygenic process, with natural selection sorting a large pool of standing genetic variation involving many adaptive alleles each resulting in environment-dependent trait optima (refs). Such polygenic processes allow the same trait optimum to be achievable by numerous SNPs. If DNA methylation is quantitative, then differentially methylated cytosines should be linked to numerous SNPs. While this genomic redundancy challenges the detection of causal SNP-DMC associations, it allows individuals with distinct genetic backgrounds to evolve to similar environmental stressors. Here, we find stress-induced SNP-DMC co-variation between several drought-responsive DMCs and hundreds of SNPs, suggesting that methylation levels at stress-responsive cytosines can be regulated via a multitude of pathways. The complete absence of covariation with SNPs of DMCs resulting from ecological divergence along a parental fine-scale gradient, on the other hand, indicates that inherited DMCs are uncoupled from most genetic variation. This suggests that (i) stress-sensitive DNA methylation is a highly quantitative trait that can be regulated by many SNPs and (ii) natural selection reduces redundancy in adaptive genetic variation associated with epigenetic variation, consequently resulting in undetectable SNP-DMC covariation. An alternative explanation for the lack of SNP-DMC co-variation is that DMCs can act as gene-independent targets of selection, but the observation of high stress-induced DMC-SNP covariation is in favour of polygenic regulation and adaptation.

We also found that the majority of DMCs were genomically separated from covarying SNPs. Although this finding was somewhat surprising, *trans*-regulation of genes appears to be relatively common and in support of omnigenic trait regulation^45,46^. While the genomic distance between DMCs and SNPs was a poor predictor of DMC-SNP covariation, DMC density appears key to the degree of covariation between SNPs and DMCs. Specifically, solitary DMCs were significantly more likely to co-vary with SNPs than DMCs clustered into islands, but only for background SNPs. The observation that outlier SNPs and DMC islands are not involved in detectable SNP-DMC covariation corroborates the notion that the process of natural selection reduced detectable SNP-DMC covariation. In this respect, DMC islands, which have been associated with important phenotypic effects, may be more important targets of natural selection than solitary DMCs.

In the framework of polygenic adaptation shaping genome-wide methylation levels, reduced genomic redundancy underlying quantitative traits under selection is expected to reduce SNP-DMC covariation in genes underlying adaptive trait variation. We correspondingly found no SNP-DMC covariation in the vicinity of flowering genes. Although methylation profiling of leaf tissue prevented us from capturing most epigenetic variation in flowering genes, we did find cytosines with significantly differentiated methylation levels in flowering genes, and none of these flowering DMCs co-varied with SNPs in flowering genes. Considering that flowering is a highly complex trait that is under strong selection along the fine-scale gradient studied here, the lack of SNP-DMC covariation across flowering genes supports a role for adaptation in uncoupling epigenetic from genetic variation.

While cross-study system experimental validation of the framework is required for making general statements regarding the role of genetic variation in driving genome-wide methylation patterns, our findings indicate that (i) DNA methylation is a quantitative trait rather than an independent evolutionary force, and (ii) DNA methylation profiling allows detecting signatures of adaptive evolution that are sensitive to genomic redundancy and genomic parallelism. Specifically, while gene-environment association studies allow detecting SNPs that align with environmental variables, they are unable to detect adaptive SNPs underlying complex traits due to the possibility of many alternative SNP-environment associations across the same environmental gradient. Because part of the genomewide DNA methylation is directly linked to phenotypic expression under selection, irrespective of the genetic background, DNA methylation scans are more likely than SNP scans to capture genome-wide levels of adaptation and trait heritability.

The amount of SNP-DMC covariation may be conservative, mainly because the bisulfite treatment of our genomes reduces the detectability of many SNPs. However, this did not prevent us from finding notable differences in SNP-DMC covariation for stress-induced DMCs as compared to inherited, drought stress-insensitive DMCs. An increased SNP calling resolution may have, at best, provided a few SNPs co-varying with inherited DMCs, but it is unlikely that this would change the overall conclusion that SNP-DMC covariation is much stronger for stress-induced DMCs than for inherited DMCs.

## ACKNOWLEDGMENTS

Hanne De Kort holds an FWO (Flemish Research Foundation) scholarship (12P6521N).

## AUTHOR CONTRIBUTIONS

Hanne De Kort developed the hypotheses, performed the sampling and analyses, and wrote the first version of this manuscript. Tuomas Toivainen performed the SNP calling. All authors provided feedback for improvement.

## REFERENCES

1. Fisher, R. A. The genetical theory of natural selection. Dover Publications, Toronto, ON. (1957).

2. Wright, S. On the Roles of Directed and Random Changes in Gene Frequency in the Genetics of Populations. Evolution (N. Y). 2, 279 (1948).

3. Payne, J. L. & Wagner, A. The causes of evolvability and their evolution. Nature Reviews Genetics 20, 24–38 (2019).

4. Meröndun, J., Murray, D. L. & Shafer, A. B. A. Genome-scale sampling /suggests cryptic epigenetic structuring and insular divergence in Canada lynx. Mol. Ecol. 28, mec.15131 (2019).

5. Wogan, G. O. U., Yuan, M. L., Mahler, D. L. & Wang, I. J. Genome-wide epigenetic isolation by environment in a widespread Anolis lizard. Mol. Ecol. 29, 40–55 (2020).

6. De Kort, H., Panis, B., Deforce, D., Van Nieuwerburgh, F. & Honnay, O. Ecological divergence of wild strawberry DNA methylation patterns at distinct spatial scales. Mol. Ecol. 29, 4871–4881 (2020).

7. Cavalli, G. & Heard, E. Advances in epigenetics link genetics to the environment and disease. Nature 571, 489–499 (2019).

8. Verhoeven, K. J. F., Jansen, J. J., van Dijk, P. J. & Biere, A. Stress-induced DNA methylation changes and their heritability in asexual dandelions. New Phytol. 185, 1108–18 (2010).

9. Herman, J. J. & Sultan, S. E. DNA methylation mediates genetic variation for adaptive transgenerational plasticity. Proc. R. Soc. B Biol. Sci. 283, 20160988 (2016).

10. Berbel-Filho, W. M., Rodríguez-Barreto, D., Berry, N., Garcia De Leaniz, C. & Consuegra, S. Contrasting DNA methylation responses of inbred fish lines to different rearing environments. Epigenetics 14, 939–948 (2019).

11. Xu, J. et al. Population-level analysis reveals the widespread occurrence and phenotypic consequence of DNA methylation variation not tagged by genetic variation in maize. Genome Biol. 20, 1–16 (2019).

12. Yagound, B., Smith, N. M. A., Buchmann, G., Oldroyd, B. P. & Remnant, E. J. Unique DNA Methylation Profiles Are Associated with cis-Variation in Honey Bees. Genome Biol. Evol. 11, 2517–2530 (2019).

13. Herrera, C. M. & Bazaga, P. Epigenetic differentiation and relationship to adaptive genetic divergence in discrete populations of the violet Viola cazorlensis. New Phytol. 187, 867–76 (2010).

14. Schmid, M. W. et al. Contribution of epigenetic variation to adaptation in Arabidopsis. Nat. Commun. 9, 4446 (2018).

15. Stajic, D., Perfeito, L. & Jansen, L. E. T. Epigenetic gene silencing alters the mechanisms and rate of evolutionary adaptation. Nat. Ecol. Evol. 3, 491–498 (2019).

16. Kankel, M. W. et al. Arabidopsis MET1 cytosine methyltransferase mutants. Genetics 163, 1109–1122 (2003).

17. Wagner, G. P. & Zhang, J. The pleiotropic structure of the genotype-phenotype map: the evolvability of complex organisms. Nat. Rev. Genet. 12, 204–13 (2011).

18. Smith, G., Smith, C., Kenny, J. G., Chaudhuri, R. R. & Ritchie, M. G. Genome-Wide DNA Methylation Patterns in Wild Samples of Two Morphotypes of Threespine Stickleback (Gasterosteus aculeatus). Mol. Biol. Evol. 32, 888–95 (2015).

19. Frachon, L. et al. Intermediate degrees of synergistic pleiotropy drive adaptive evolution in ecological time. Nat. Ecol. Evol. 1, 1551–1561 (2017).

20. Csilléry, K., Rodríguez-Verdugo, A., Rellstab, C. & Guillaume, F. Detecting the genomic signal of polygenic adaptation and the role of epistasis in evolution. Mol. Ecol. 27, 606–612 (2018).

21. Gugger, P. F., Fitz-Gibbon, S., PellEgrini, M. & Sork, V. L. Species-wide patterns of DNA methylation variation in *Quercus lobata* and their association with climate gradients. Mol. Ecol. 25, 1665–1680 (2016).

22. Wang, M. Z., Li, H. L., Li, J. M. & Yu, F. H. Correlations between genetic, epigenetic and phenotypic variation of an introduced clonal herb. Heredity (Edinb). 124, 146–155 (2020).

23. Gáspár, B., Bossdorf, O. & Durka, W. Structure, stability and ecological significance of natural epigenetic variation: a large-scale survey in Plantago lanceolata. New Phytol. 221, 1585–1596 (2019).

24. Cortijo, S. et al. Mapping the epigenetic basis of complex traits. Science 343, 1145–8 (2014).

25. Jeong, H., Wu, X., Smith, B. & Yi, S. V. Genomic landscape of methylation islands in hymenopteran insects. Genome Biol. Evol. 10, 2766–2776 (2018).

26. Jeziorska, D. M. et al. DNA methylation of intragenic CpG islands depends on their transcriptional activity during differentiation and disease. Proc. Natl. Acad. Sci. U. S. A. 114, E7526–E7535 (2017).

27. Suzuki, M. M. & Bird, A. DNA methylation landscapes: provocative insights from epigenomics. Nat. Rev. Genet. 9, 465–76 (2008).

28. Gent, J. I. et al. CHH islands: De novo DNA methylation in near-gene chromatin regulation in maize. Genome Res. 23, 628–637 (2013).

29. Zhang, H., Lang, Z. & Zhu, J. K. Dynamics and function of DNA methylation in plants. Nature Reviews Molecular Cell Biology 19, 489–506 (2018).

30. Kawakatsu, T. et al. Unique cell-type-specific patterns of DNA methylation in the root meristem. Nat. Plants 2, 16058 (2016).

31. Dapp, M. et al. Heterosis and inbreeding depression of epigenetic Arabidopsis hybrids. Nat. Plants 1, 15092 (2015).

32. Latzel, V., Rendina González, A. P. & Rosenthal, J. Epigenetic Memory as a Basis for Intelligent Behavior in Clonal Plants. Front. Plant Sci. 7, 1354 (2016).

33. Ardura, A., Zaiko, A., Morán, P., Planes, S. & Garcia-Vazquez, E. Epigenetic signatures of invasive status in populations of marine invertebrates. Sci. Rep. 7, 42193 (2017).

34. Artemov, A. V. et al. Genome-Wide DNA Methylation Profiling Reveals Epigenetic Adaptation of Stickleback to Marine and Freshwater Conditions. Mol. Biol. Evol. 34, 2203–2213 (2017).

35. Thorson, J. L. M. et al. Epigenetics and adaptive phenotypic variation between habitats in an asexual snail. Sci. Rep. 7, 14139 (2017).

36. Rendina González, A. P., Preite, V., Verhoeven, K. J. F. & Latzel, V. Transgenerational Effects and Epigenetic Memory in the Clonal Plant Trifolium repens. Front. Plant Sci. 9, 1677 (2018).

37. Wibowo, A. et al. Partial maintenance of organ-specific epigenetic marks during plant asexual reproduction leads to heritable phenotypic variation. Proc. Natl. Acad. Sci. 115, E9145–E9152 (2018).

38. De Kort, H. et al. Pre-adaptation to climate change through topography-driven phenotypic plasticity. J. Ecol. 108, 1465–1474 (2020).

39. Andrés, F. & Coupland, G. The genetic basis of flowering responses to seasonal cues. Nature Reviews Genetics 13, 627–639 (2012).

40. Bratzel, F. & Turck, F. Molecular memories in the regulation of seasonal flowering: From competence to cessation. Genome Biology 16, 1–14 (2015).

41. Shulaev, V. et al. The genome of woodland strawberry (Fragaria vesca). Nat. Genet. 43, 109–116 (2011).

42. Akalin, A. et al. methylKit: a comprehensive R package for the analysis of genome-wide DNA methylation profiles. Genome Biol. 13, R87 (2012).

43. Luu, K., Bazin, E. & Blum, M. G. B. pcadapt: an R package to perform genome scans for selection based on principal component analysis. in Molecular Ecology Resources 17, 67–77 (Blackwell Publishing Ltd, 2017).

44. Dray, S., Dufour, A., Thioulouse, J., Jombart, T. & Pavoine, S. Ade4: analysis of ecological data: exploratory and euclidean methods in environmental sciences. R Packag. version (2009).

45. Boyle, E. A., Li, Y. I. & Pritchard, J. K. An Expanded View of Complex Traits: From Polygenic to Omnigenic. Cell 169, 1177–1186 (2017).

46. Barghi, N., Hermisson, J. & Schlötterer, C. Polygenic adaptation: a unifying framework to understand positive selection. Nat. Rev. Genet. 1–13 (2020). doi:10.1038/s41576-020-0250-z

